# Integrated pathogen load and dual transcriptome analysis of systemic host-pathogen interactions in severe malaria

**DOI:** 10.1101/193631

**Authors:** Hyun Jae Lee, Michael Walther, Athina Georgiadou, Davis Nwakanma, Lindsay B. Stewart, Michael Levin, Thomas D. Otto, David J. Conway, Lachlan J. Coin, Aubrey J. Cunnington

**Author notes:** Corresponding author (AJC). Denotes equal contribution (Thomas D. Otto, David J. Conway, Lachlan J. Coin).

## Abstract

The pathogenesis of severe *Plasmodium falciparum* malaria is incompletely understood. Since the pathogenic stage of the parasite is restricted to blood, dual RNA-sequencing of host and parasite transcripts in blood can reveal their interactions at a systemic scale. Here we identify human and parasite gene expression associated with severe disease features in Gambian children. Differences in parasite load explained up to 99% of differential expression of human genes but only a third of the differential expression of parasite genes. Co-expression analyses showed a remarkable co-regulation of host and parasite genes controlling translation, and host granulopoiesis genes uniquely co-regulated and differentially expressed in severe malaria. Our results indicate that high parasite load is the proximal stimulus for severe *P. falciparum* malaria, that there is an unappreciated role for many parasite genes in determining virulence, and hint at a molecular arms-race between host and parasite to synthesise protein products.

## Introduction

*Plasmodium falciparum* malaria is one of the most important infectious diseases affecting humankind[1]. Progress has been made in malaria treatment and control in the last decade but this is threatened by the spread of antimalarial and insecticide resistance[2-4] Understanding of pathogenic mechanisms associated with severe malaria (SM), which puts individuals at risk of death, has also progressed[1, 5, 6]. Immunopathology, vascular endothelial dysfunction and parasite sequestration (obstruction of the microvasculature by cytoadherent parasites) all have putative roles in SM[5], and high parasite load is also strongly associated with greater risk of severe disease[5, 7-10]. Rodent models have contributed to mechanistic dissection of the pathogenic processes, but these cannot yet reproduce all of the features of naturally-occurring *P. falciparum* malaria[6, 11]. An integrated understanding of the respective roles and interactions of host and parasite in human SM is notably lacking, and whether SM involves excessive, proportionate or insufficient host responses to the parasite is largely unknown. Here we combine estimates of parasite load with host and parasite whole blood gene expression to investigate their associations with severity and different pathological features of SM, aiming to provide a global view of systemic host-parasite interaction. This approach allows us to harness the natural variation which occurs between humans infected with *P. falciparum* to better understand the different pathogenic processes which underlie SM.

## Results

We performed dual-RNA sequencing on whole blood of 46 Gambian children with uncomplicated (UM, n=21) and severe (SM, n=25) *P. falciparum* malaria (Supplementary Table 1). After exclusion of *var*, *rifin*, *stevor[12]* and highly polymorphic regions in the parasite genome (see Methods), we obtained median 26.6 million (26.6 million SM, 26.7 million UM, P=0.913) human and 9.61 million (10.3 million SM, 5.03 million UM, P=0.346) parasite uniquely mapped reads from each subject (Fig 1a), with considerably greater parasite read depth than a previous study conducted in adults with UM[13]. Systemic infection provokes changes in blood leukocyte subpopulations which could dominate changes in gene expression[14] so we performed “gene signature” based deconvolution[15] to identify and adjust for heterogeneity in the major leukocyte subpopulations in each sample (Fig 1b, Supplementary Fig 1). Parasite gene expression *in vivo* is also influenced by the mixture of parasite developmental stages at the time of sampling because there is phasic variation in gene expression[16] and increasing RNA content during the intraerythrocytic developmental cycle[17]. Therefore we used the same deconvolution approach with “gene signatures” derived from highly synchronous parasite cultures[16, 18] to identify the contribution of parasites at different developmental stages (Supplementary Fig 2 and Fig 1c). There was a trend towards greater proportions of late stage asexual parasites and gametocytes in children with SM (Fig 1d).

**Figure 1.**
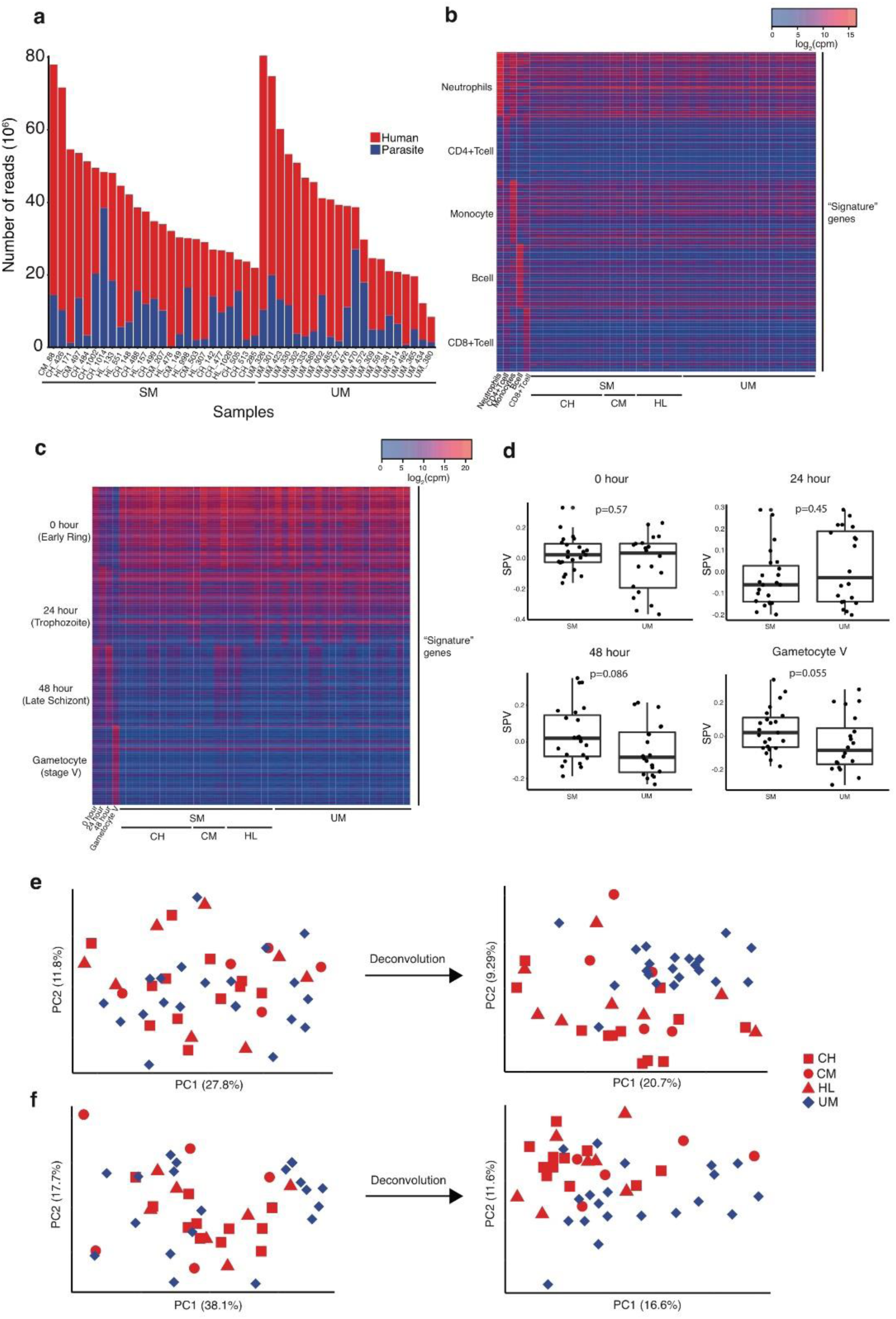
Whole blood dual RNA-sequencing and deconvolution. (**a**) Uniquely mapped reads from human (red) and *P. falciparum* (blue) from subjects with severe (SM, n=25) and uncomplicated malaria (UM, n=21). (**b,c**) Heatmaps showing signature gene expression for different leukocyte (**b**) and parasite developmental stage (**c**) populations (rows) and their relative intensity in individual subjects with SM, including different SM phenotypes (CH, cerebral malaria plus hyperlactatemia; CM, cerebral malaria; HL, hyperlactatemia), and UM (columns). (**d**) Surrogate proportion variables for parasite developmental stages compared between SM and UM using the Mann-Whitney test (bold line, box and whiskers indicate median, interquartile range and 1.5-times interquartile range respectively). (**e,f**) Principal component plots showing the effect of deconvolution on the segregation of subjects with UM and SM, adjusting human (**e**) and parasite (**f**) gene expression for differences in proportions of leukocytes or parasite developmental stages respectively. Analyses of human gene expression (**b**,**e**): SM, n=25; UM, n=21. Analyses of parasite gene expression (**c**,**d**, **f**): SM, n=23; UM, n=20.

Examination of principal component plots before and after adjustment for heterogeneity in the mixture of leukocytes and parasite developmental stages revealed that segregation of SM and UM cases was improved after adjustment (Fig 1e,f). Therefore we used these adjusted gene expression values for all subsequent analyses, essentially allowing us to compare gene expression as if all subjects had the same leukocyte and parasite population compositions.

Whole blood genome-wide gene expression can be used to characterise host cellular responses and infer upstream regulators[19], whilst variations in *P. falciparum* gene expression are believed to reflect adaptation to the host environment and contribute to virulence[20, 21]. We identified significantly differentially expressed genes from human and parasite in SM vs UM (Fig 2a,b) and also for different subtypes of SM (hyperlactatemia (HL) and cerebral malaria (CM), alone or in combination) vs UM (Supplementary Figure 3). There were 770 human and 236 parasite significantly differentially expressed genes (DEGs, with false discovery rate (FDR)-adjusted P<0.05) between SM and UM (Supplementary Table 2 and 3). Some human genes had both conspicuously high expression in SM relative to UM (high log-fold change) and were also highly significant (Fig 2a): the four most upregulated (*MMP8*, matrix metallopeptidase 8; *OLFM4*, olfactomedin 4; *DEFA3*, defensin A3; *ELANE*, neutrophil elastase) notably all encode neutrophil granule proteins[22]. Interestingly the number of human and parasite DEGs was substantially higher when comparing the subgroup of subjects with cerebral malaria plus hyperlactatemia (CH, the most severe phenotype, n=12) vs UM, despite the smaller number of subjects (Supplementary Table 2, Supplementary Figure 3).

**Figure 2.**
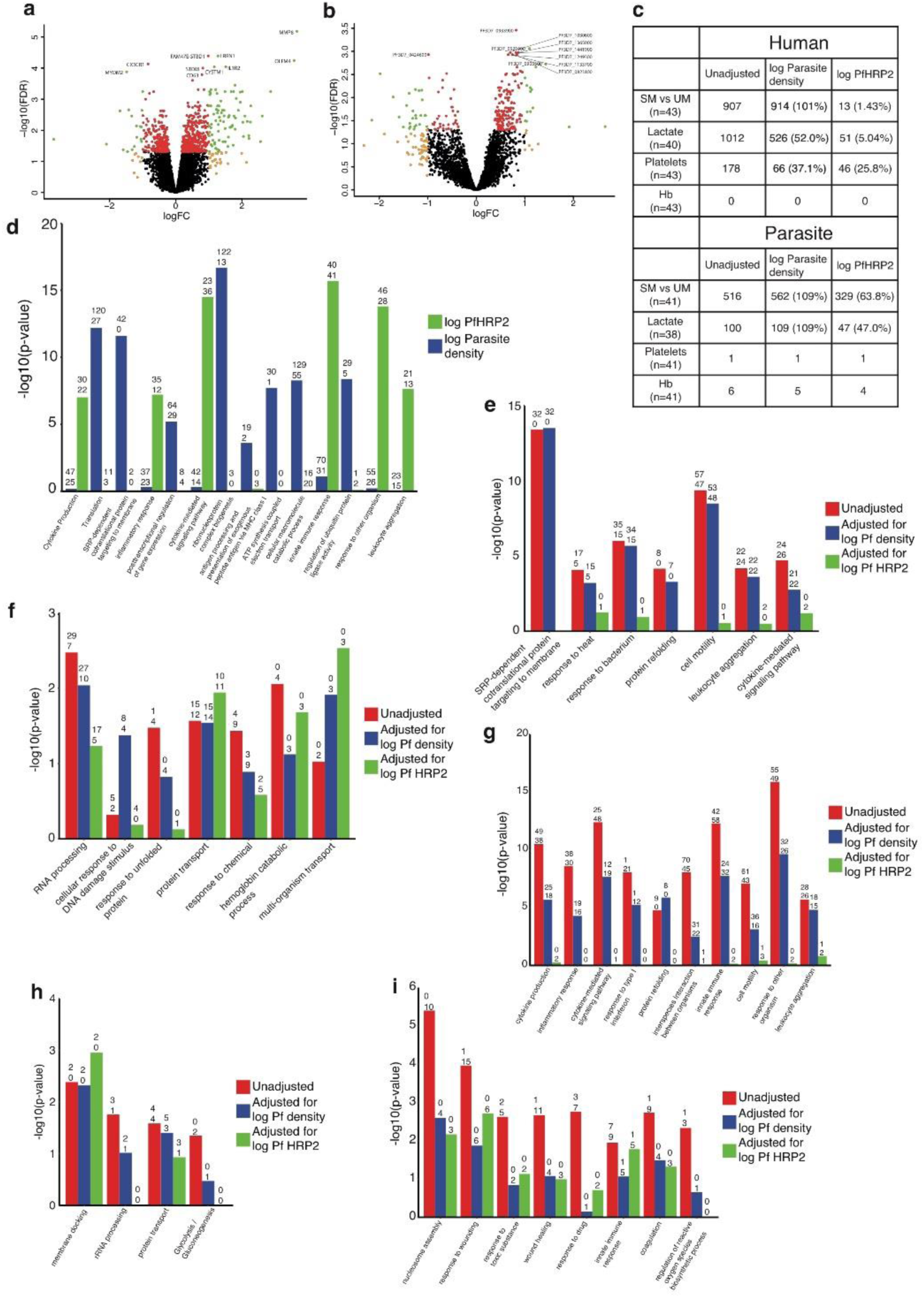
Association of gene expression with severity features and dependency on parasite load. (**a, b**) Volcano plots showing extent and significance of up- or down- regulation of human (**a**) or *P. falciparum* (**b**) gene expression in SM compared with UM (red and green, P <0.05 after Benjamini-Hochberg adjustment for false discovery rate (FDR); orange and green, absolute log2-fold change (FC) in expression > 1; the 10 most significant genes are annotated; human comparison SM n=25, UM=21; parasite comparison SM n=23, UM=20). (**c**) Number of human and parasite genes associated with severity category and laboratory markers of severity before and after adjustment for parasite load expressed as either log circulating parasite density or log PfHRP2 concentration. Only subjects with complete data for every parameter are included. (**d-i**) Most significantly enriched, non-redundant, gene ontology terms for genes significantly associated with log parasite density (**d**) and log PfHRP2 (**e**), and the effect of adjustment for these measures on genes significantly associated with severity (**f-i**) (numbers above bars indicate the number of up-regulated/positively-associated and down-regulated/negatively-associated genes within each category). (**e,f**) Human (**e**) and *P. falciparum* (**f**) genes significantly differentially expressed in SM vs UM. (**g,h**) Human (**g**) and *P. falciparum* (**h**) genes significantly correlated with blood lactate concentration. (**i**) Human genes significantly correlated with platelet count.

Previous studies have shown a correlation between host gene expression and circulating parasitemia[23, 24], suggesting that this might explain some of the differences in gene expression between SM and UM. However peripheral blood parasite measurements underestimate the total number of parasites in the body because parasitized red blood cells can also become sequestered, accumulating in small blood vessels rather than remaining in circulation[12, 25]. The parasite protein, *P. falciparum* histidine rich protein 2 (PfHRP2), can be used as a plasma biomarker of total parasite load and is more strongly associated with severity[7, 8, 10] (Supplementary Table 1) and death[8, 10]. Therefore we examined the association of host and parasite gene expression with both circulating parasite density and PfHRP2 (restricting comparisons to subjects with data for both). We found 1886 human genes significantly (FDR P<0.05) correlated with log parasite density and 616 significantly correlated with log PfHRP2 (102 common to both), whilst only 2 and 10 parasite genes were significant in the corresponding analyses (none common to both) (Supplementary Tables 4 and 5). We then asked to what extent the differences between SM and UM phenotypes were dependent on parasite load. The number of human SM vs UM DEGs remained almost unchanged before and after adjustment for parasite density but was reduced by 98.6% after adjustment for PfHRP2, whilst parasite DEGs changed much less after the same adjustments (Figure 2c, Supplementary Tables 2 and 3). Findings were similar when adjusting for parasite load in comparisons of each of the SM subtypes vs UM (Supplementary Tables 2 and 3).

Genes associated with severity after adjustment for parasite load may include determinants of susceptibility to severe disease. Of particular interest amongst these, MMP8 (also known as collagenase 1) is a metallopeptidase which causes endothelial barrier damage in several infection models[26, 27]; *AZI2* (also known as NF-Kappa-B-Activating Kinase-Associated Protein 1, NAP1) encodes a regulator of the type 1 interferon response[28], a pathway which is known to control severity of disease in rodent malaria models[29]; whilst CX3CR1 is the receptor for fractalkine (a biomarker of CM in humans[30]) and a marker for a subset of monocytes which are particularly efficient at killing malaria parasites[31].

Next we performed pathway analyses to better understand the biological functions of the significant genes in the preceding analyses. Human genes correlated with log parasite density were particularly enriched in pathways related to translation (especially exported proteins), oxidative phosphorylation and ubiquitination (Fig 2d, Supplementary Table 6), with predicted upstream regulation by RICTOR (RPTOR independent companion of MTOR complex 2), HNF4A (hepatocyte nuclear factor 4 alpha) and XBP1 (X-box binding protein 1; Supplementary Table 7). Genes correlated with log PfHRP2 were particularly enriched in inflammatory and immune response functions (specifically innate response and type 1 interferon, Fig 2d, Supplementary Table 6), with predicted regulation by IFN-γ, TGM2 (transglutaminase 2) and IFN-α2 (Supplementary Table 7). Some of these immune response functions were also correlated with parasite density but were not significantly enriched because of the larger denominator in this analysis (Fig 2d, Supplementary Tables 6). We compared human DEGs from SM vs UM comparisons without adjustment and with adjustment for parasite density or PfHRP2 (Figure 2e, Supplementary Table 6)). In unadjusted analyses human genes were particularly enriched in processes controlling protein synthesis and targeting to the endoplasmic reticulum, cell stress and immune response and the most significant predicted upstream regulators were CSF3 (colony stimulating factor 3, also known as granulocyte colony stimulating factor, GCSF), FAS (Fas cell surface death receptor) and PTGER2 (Prostaglandin E receptor 2, Supplementary Table 7). After adjustment for parasite density we observed little change in pathway enrichment associated with severe malaria, whilst adjustment for PfHRP2 reduced all pathway enrichments. In contrast parasite pathways were less influenced by adjustment for either parasite density or PfHRP2 (Fig 2f, Supplementary Table 8), the most consistently significant being RNA processing, protein transport, and hemoglobin catabolism. Findings were broadly similar in corresponding analyses for all subtypes of SM (Supplementary Figure 4, Supplementary Table 6), with the exception that adjustment for parasite density produced a 5-fold increase in the number of DEGs in the hyperlactatemia (HL) vs UM comparison. This intriguing finding suggests that the response to circulating parasites in these groups with similar parasite density (Supplementary Table 1) may partially mask differences in gene expression associated with hyperlactatemia.

To further explore pathophysiology we examined the correlation of human and parasite gene expression with lactate and hemoglobin concentrations and platelet count[32]. 1012 human genes were significantly correlated with lactate concentration, reducing by half after adjustment for parasite density and by 95% after adjustment for PfHRP2 (Fig 2c,g, Supplementary Table 4). Immune response pathways were prominent in unadjusted analysis (the negative association with type 1 interferon being particularly notable), and the most significant predicted upstream regulators were interferon-γ, interferon-α, and TNF. Adjustment for parasite density retained most enrichment terms whilst adjustment for PfHRP2 removed almost all significant enrichment (Fig 2g, Supplementary Table 6) but remaining genes included *PKM* (encoding the glycolytic enzyme pyruvate kinase M) and *GYS1* (encoding the glycogenic enzyme glycogen synthase 1) (Supplementary Table 4). These findings suggest hyperlactatemia is driven by the parasite load-dependent inflammatory response, but also influenced by some parasite load-independent variation in control of host metabolism. 100 parasite genes were significantly correlated with lactate, with much less dependency on parasite load (Fig 2c,h, Supplementary Table 5). Unexpectedly these included two glycolysis genes (hexokinase and acetylCoA synthetase), negatively correlated with lactate (Fig 2h, Supplementary Table 9), suggesting that rather than parasite-derived lactate driving hyperlactatemia, host-derived lactate may negatively regulate parasite glycolysis. Compared to lactate, platelet count was associated with fewer human genes, and these were less dependent on parasite load (Fig 2c,i, Supplementary Table 4). The most enriched pathways also differed considerably, with nucleosome assembly (predominantly histone genes), coagulation, and response to wounding genes, all negatively correlated with platelet count (Fig 2i), and the most significant predicted upstream regulators being IL13, RB1 (RB transcriptional corepressor 1), and IL1RN (Supplementary Table 7). Activation of coagulation pathways is increasingly recognised in severe malaria[33, 34], but free histones can also induce thrombocytopenia[35] and may be relevant in malaria. No human genes and few parasite genes were associated hemoglobin concentration in any analyses and only one parasite gene was associated with platelet count (Figure 2c, Supplementary Table 4,5).

Taken together the preceding findings indicate that total parasite load is the dominant driver of host leukocyte gene expression in malaria, particularly inflammatory and immune response genes, and differences in parasite load explain almost all of the human gene expression differences between SM and UM. Despite this, specific parasite load-dependent pathways were differentially associated with distinct aspects of systemic pathophysiology, and circulating parasites correlated with patterns of host gene expression suggesting that parasite localization substantially alters the host-parasite interaction. In contrast to host genes, parasite gene expression showed little association with parasite load, implying that the non-polymorphic genes differentially expressed between SM and UM do not directly contribute to high parasite load, but may contribute to other aspects of pathogenesis or simply reflect parasite responses to the perturbed host environment.

Whilst these independent analyses of host and parasite gene expression associations with severity are enlightening, dual-RNA sequencing can also be used to identify molecular interactions within and between species and their associations with severity[36]. Expression of groups of genes with common functional roles are often highly correlated and can be identified through co-expression network analysis[37]. We applied this methodology to identify correlated modules of genes, which could originate from either or both species and were named according to the “hub gene” which has the greatest connectivity within the module. First we analysed all subjects together and generated a network with 26 modules (Fig 3 and Supplementary Table 9): 10 containing exclusively human genes, 5 exclusively parasite genes, and 11 with both human and parasite genes (although most of these were highly skewed to a single species). All modules showed significant functional enrichments regardless of host or parasite origin. The composite expression of genes within a module can be described by a module eigengene value and, as expected, there were significant associations between module eigengene values and severity, parasite load, and laboratory parameters (Figure 3). Only the *HSPH1* (heat shock protein family H (Hsp110) member 1) module contained more than 10 genes from both human and parasite, strongly enriched in human heat shock response genes and parasite RNA metabolism genes, perhaps indicating that these parasite genes particularly promote human cell stress. Some host-dominated and parasite-dominated modules were also highly correlated with each other, most notably the *RPL24* (ribosomal protein L24) module (highly enriched in translation pathways) was strongly correlated with the remarkably homologous *PF3D7_0721600* (putative 40S ribosomal protein S5) parasite module. We excluded read mapping errors as an explanation for this, and suggest that this indicates co-regulation of conserved host and parasite translation machinery. Furthermore, most of these genes were also differentially expressed between SM and UM, perhaps indicating a “molecular arms race” between parasite and host to synthesise proteins which may, in excess, contribute to collateral tissue damage.

**Figure 3.**
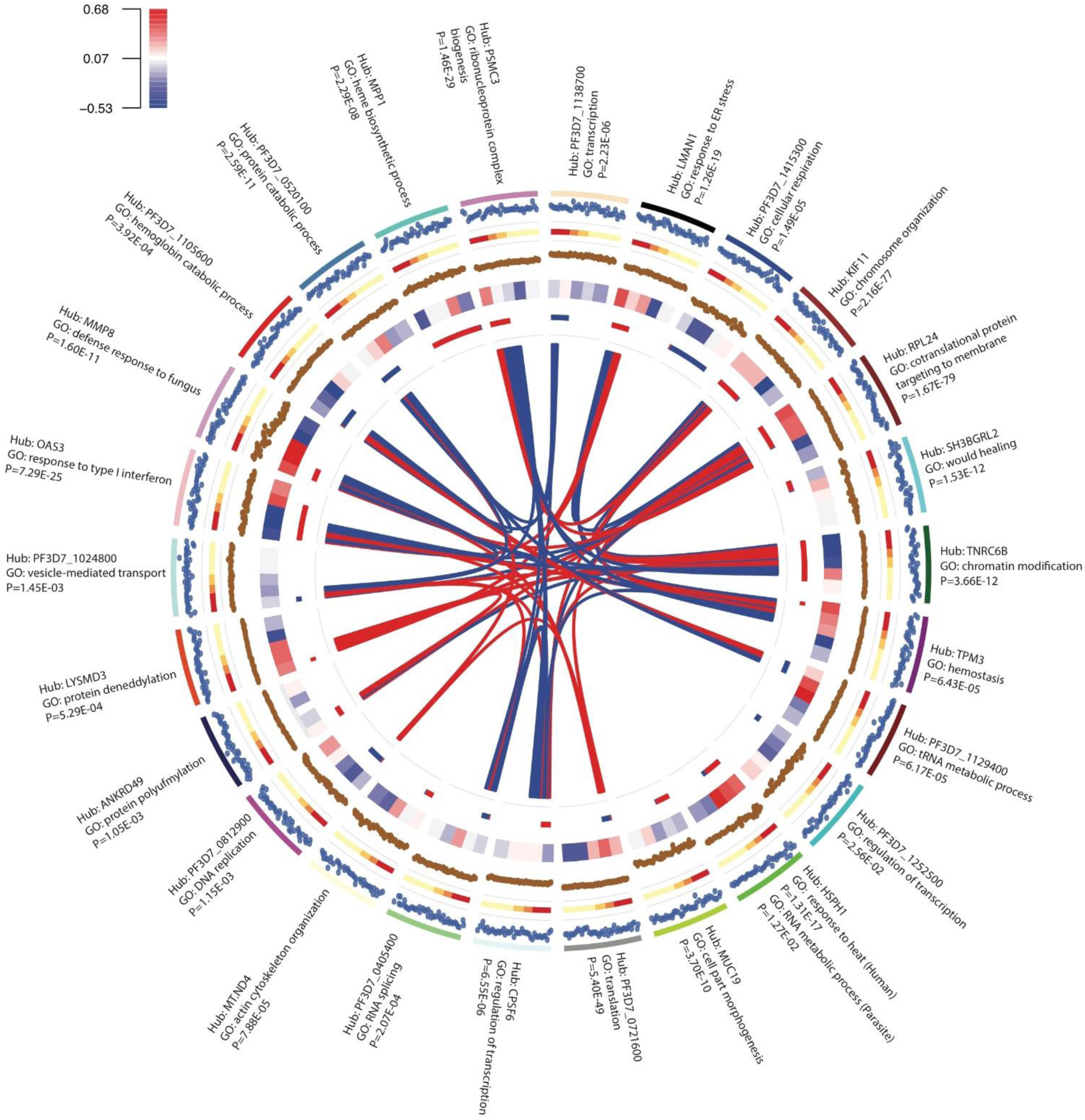
Interspecies gene expression modules and their associations with severity. Circos plot showing gene expression modules obtained from whole genome correlation network analysis using expression of all human and parasite genes from each subject (SM, n=22; UM, n=19) as the input. From outside to inside: labels, hub gene and most enriched GO term (with enrichment P-value) for each module; track 1, module eigengene value for each subject; track 2, clinical phenotype (Red=CH, Orange=CM, Green=HL, Yellow=UM); track 3, hub gene expression (log CPM) for each subject; track 4, heatmap for correlation with laboratory measurements (clockwise: log parasite density, log PfHRP2, lactate, platelets, haemoglobin; colour intensity represents Pearson correlation coefficient as shown in legend); track 5, module size and composition (length proportional to number of genes in module; red, human genes; blue, parasite genes); polygons connect modules with significant (FDR P<0.01) Pearson correlation between eigengene values (width proportional to −log10 FDR P-value; red=positive correlation, blue=negative correlation)

Co-expression network modules can be used as units of analysis, affording considerable dimension reduction for whole-genome expression data. We used module eigengene values[37, 38] and parasite load (with which many modules were correlated, Fig 3) in linear regression models to determine the best within-sample predictors of severity, starting with all significant univariate associations and proceeding by backward selection (Supplementary Table 10). The best multivariate model combined *MMP8*, *OAS1* (2’-5’-oligoadenylate synthetase 1) and *LYSMD3* (LysM, putative peptidoglycan-binding, domain containing 3) module eigengenes, but not parasite load. Interestingly these modules represent distinct aspects of the immune response: the *MMP8* module, highly enriched in defence response genes with predicted upstream regulators CEBPA (CCAAT/enhancer binding protein alpha, a myeloid transcription factor) and CSF3, likely reflects granulopoiesis[22]; the *OAS1* module is highly enriched for type 1 interferon response genes; the small *LYSMD3* module, with limited GO enrichment, contains a functional network around interferon-γ (Supplementary Figure 5). The direction of association of the *OAS1* module with severity changed from negative in univariate analysis to positive in the multivariate analysis, suggesting that inadequate downregulation of the type-1 interferon response in conjunction with upregulation of granulopoiesis and interferon-γ signalling may contribute to pathogenesis.

Considering all subjects together for generation of co-expression networks maximises power to detect consistently co-regulated genes but may not identify sets of genes where co-regulation is altered by severity. For this reason we also created separate co-expression networks for UM and SM and compared the modules to identify differential co-expression (Fig 4, Supplementary Table 11). Eight modules showed significant preservation between networks, seven were partially preserved, and two were unique to SM (Figure 4a, Supplementary Table 11). Partial preservation was common amongst modules comprised predominantly from human or parasite genes (Figure 4a,b), and module preservation was not dependent on the proportion of module genes differentially expressed between SM and UM (Figure 4a,c). Again, a *MMP8* module was identified (exclusively human genes, many encoding neutrophil granule and phagosome components), unique to SM with 38% of genes significantly differentially expressed between SM and UM, enriched in host defence pathways (Supplementary Table 11) and predicted to be regulated by CEBPA, CSF3 and TNF. These findings strongly suggest this module represents emergency granulopoiesis[22] and mark this as a specific feature of SM. The *TIPRL* (TOR Signaling Pathway Regulator) module (99.2% human genes) was also unique to SM but contained very few (1.3%) DEGs, had limited GO enrichment (Supplementary Table 11), and the most significant predicted upstream regulator was the transcription factor HNF4A. Both TIPRL and HNF4A have regulatory roles in metabolic, inflammatory and apoptosis signal pathways, so the minimal change in expression of this module may represent an aberrant response in SM. Amongst the partially preserved modules we found evidence that host and parasite translation pathways were more tightly co-regulated in SM than UM, genes being distributed across fewer modules in SM (Fig 4a, Supplementary Table 11).

**Figure 4.**
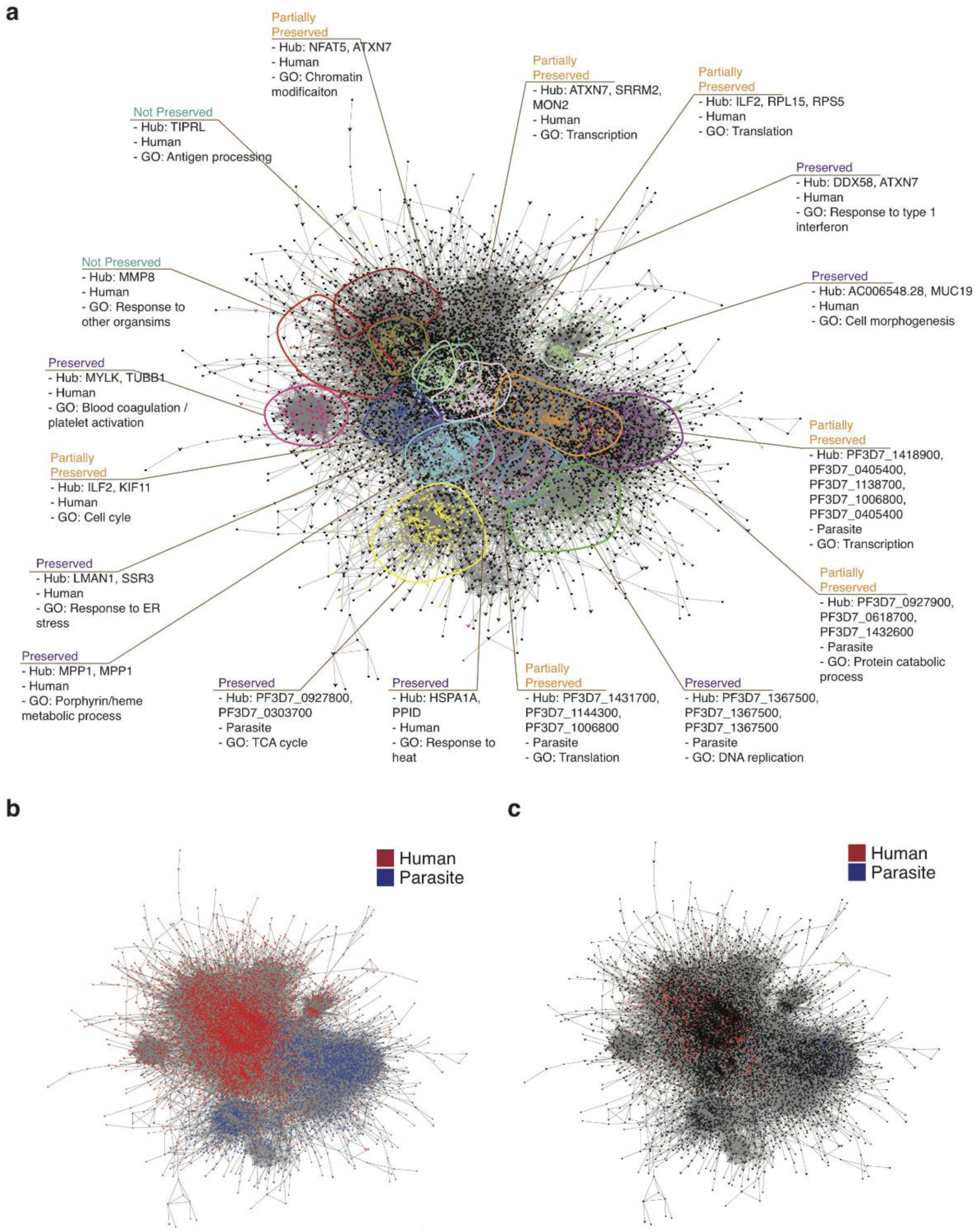
Severity-associated differential co-expression within the interspecies gene expression network.(**a-c**) Cytoscape visualisation of merged co-expression networks derived separately from SM (n=22) and UM (n=19). Networks were merged such that genes found in both sub-networks (represented as arrow-shaped, larger-sized nodes) are connected to genes found in only one sub-network (represented as circular-shaped and smaller-sized nodes). (**a**) Genes and gene clusters are coloured and annotated by module, species, most enriched gene ontology terms, and conservation between sub-networks (preserved, module pairs from SM and UM sub-networks display highly significant overlap with each other and much less significant overlap with other modules; partially preserved, module clusters display significant overlaps with two or more modules in the other subnetwork; unique, gene clustering only found in one sub-network); genes in black do not belong to any characterized module. (**b**) Identical network layout with genes coloured by species (red, human; blue, *P. falciparum*). (**c**) Identical network layout with genes coloured by whether they are significantly differentially expressed in SM vs UM (red, human; blue, *P. falciparum*; black, not differentially expressed).

## Discussion

We have shown that dual-RNA sequencing can be used to identify systemic host-pathogen interactions and potential pathogenic mechanisms associated with severe infection in humans. The differences in human and parasite gene expression between SM and UM were much clearer after adjusting for heterogeneity of leukocyte population and parasite developmental stage. Although the importance of accounting for such variation is well recognised[14], it has rarely been done in malaria or other infectious disease transcriptomic studies.

Our most striking finding came from integrating parasite load with global gene expression, revealing the overriding effect of parasite load on the differences in human gene expression between SM and UM. Previous studies have examined the association between human gene expression and circulating parasitemia[13, 23, 24], but we found that estimation of total body parasite load was necessary to appreciate the full effect on host response. Our findings imply that SM is not the consequence of an excessive host response, but that there is an appropriate host response to an excessive pathogen load. This has important implications for other infectious disease, immunology, and pathogenesis research in humans. Total body pathogen load is much harder to measure in other infections in humans[39], yet failure to account for it may lead to misinterpretation of associations between host factors and severity or protection.

Despite the dominant effect of parasite load, we found that specific sets of genes induced by infection were associated with different pathophysiological consequences of malaria. Distinct sets of genes were correlated lactate concentration and platelet count, and associated with different clinical presentations of SM. Alternative analytical approaches repeatedly identified the association of genes expressed during neutrophil granulopoiesis (such as *MMP8*) and translation pathways with severe outcomes. There is plentiful evidence that neutrophil granule proteins are released in severe malaria[40, 41], can impair vascular endothelial functions such as barrier integrity[26, 27], and may therefore have a direct role in the pathogenesis of SM. Unfortunately neutrophil related signatures are not differentially expressed in the whole blood transcriptome of the widely used rodent experimental cerebral malaria model[42], which means that experimental testing of the role of neutrophils may be challenging.

We observed an intriguing relationship between type 1 interferon responses and severity, which may help to tie together data from previous observations in humans and animal models. A previous small study found higher expression of type-1 interferon response genes in UM than SM and suggested that this may be protective against developing SM[43]. However we found that type-1 interferon response genes were negatively correlated with parasite load, indicating that downregulation with increasing parasite load (and severity) is a more likely explanation. When we performed multivariate analyses using gene expression modules to explain severity, our results suggested that insufficient downregulation of type 1 interferons was in fact associated with severity. This would be more consistent with results in several animal models where genetic or antibody-mediated ablation of type-1 interferon signalling improves outcome[44-47]

The role of translation pathways is more speculative, but co-regulation of these genes between host and parasite, which becomes tighter in more severe disease, implies that there may be an inter-species feedback loop. Increased translation is important for production of host defence effector proteins[48] and parasite proteins which enable survival[49]. Perhaps, as parasite load increases the host response increases, the parasite produces more proteins necessary to survive, and the cycle amplifies until parasite load and host response cause host organ damage and severe disease.

In addition to translation-related genes, we also identified hundreds of other parasite genes which were associated with severe disease. Many of these genes have as yet unknown function. However the enrichment of genes involved in protein transport, for example, suggests there may be layers of control which determine parasite protein export into the host cell and the molecular host-parasite interactions which predispose to SM. Some of these aspects of parasite gene regulation may only be appreciated *in vivo*, in the parasite’s natural environment.

The data we have generated and comprehensive analyses we have performed provide a unique and valuable resource for the research community. These should be launch points for future studies using alternative approaches to assess whether the mechanisms we have implicated through gene expression do indeed play causal roles in SM and may be targets for much needed adjunctive therapies.

## Methods

### Subjects and samples

Gambian children (under 16 years old) with *P. falciparum* malaria were recruited from three periurban health centres, The MRC Gate Clinic, Brikama Health Centre, and The Jammeh Foundation for Peace Hospital, Serekunda, as part of a larger study of severe malaria[7, 50, 51]. Informed consent was obtained from the child’s parent or legal guardian for collection and subsequent use of samples. The study was approved by the Gambian Government / MRC Laboratories Joint Ethics Committee. All children underwent full clinical examination and were managed in accordance with the Gambian government guidelines. Malaria was defined by the occurrence of fever in the last 48 hours before recruitment and >5000 asexual parasites/μL in the peripheral blood. Subjects were further categorized into different severe malaria phenotypes using modified World Health Organization criteria: cerebral malaria (CM) was defined as Blantyre Coma Score (BCS) of 1 or 2, or a BCS of 3 if the motor response was 1, AND no hypoglycaemia, no rapid improvement in response to fluid resuscitation, no suspicion of meningitis; hyperlactatemia (HL), blood lactate concentration > 5mmol/L; both CM and HL (CH)[7]. At the time of presentation to the clinic, prior to any antimalarial treatment or blood transfusion, capillary blood was used for measurement of lactate and glucose concentrations and thick and thin blood films, venous blood was collected into EDTA for sickle cell screen and full blood count, PAXgene blood RNA tube (BD), and sodium heparin (BD) for plasma separation[50]. Parasitemia was calculated using 50 high power fields on Giemsa-stained thin blood smears. Plasma *P. falciparum* histidine-rich protein II (PfHRP2) was measured by ELISA (Cellabs)[7].

For the present study we used 46 subjects selected from those with ≥1μg RNA available which showed no / minimal evidence of degradation on visual inspection of a Bioanalyser (Agilent) trace (RNA integrity number calculations are not valid for dual species RNA analysis). To reduce potential confounding we aimed to frequency match subjects between SM and UM groups as closely as possible by age and gender, and if there remained a choice of samples available we selected those with the most complete additional clinical and laboratory data. For UM samples we aimed to include an equal number with parasitemia above and below 5% (to maximise the chance of obtaining parasite reads in some of the subjects). For SM samples we aimed to include subjects with each of the common SM phenotypes seen in this part of the Gambia in approximately equal numbers, although final numbers were determined by availability and quality of RNA. Detailed information about the study subjects is shown in Supplementary Table 1 and Supplementary Dataset 1. Characteristics were compared between subject groups using one-way ANOVA for continuous data and Fisher’s exact test for categorical data.

### RNA sequencing

Total RNA was extracted using the PAXgene Blood RNA kit (BD). Libraries were prepared from 1μg of total RNA using the ScriptSeq v2 RNA-seq library preparation kit (Illumina) with additional steps to remove ribosmal RNA (rRNA) and globin messenger RNA (mRNA) using the Globin-Zero Gold kit (Epicentre). Strand-specific libraries were sequenced using the 2x100 bp protocol with an Illumina HiSeq 2500 instrument. In order to eliminate batch effects, samples were randomized for the order of library preparation. For sequencing, 5-6 samples were run per lane, and each lane contained at least one sample from each disease type, randomly allocated in a block design. Library preparation and sequencing were carried out by Exeter University sequencing service.

### Genomes and RNA annotations

Human reference genome (hg38) was obtained from UCSC genome browser (http://genome.ucsc.edu/) and *P. falciparum* reference genome (release 24) was obtained from PlasmoDB (http://plasmodb.org/). Human gene annotation was obtained from GENCODE (release 22) (http://gencodegenes.org/releases/) and *P. falciparum* gene annotation from PlasmoDB (release 24) (<u>http://plasmodb.org</u>).

### Read Mapping and quantification

RNA-seq data was mapped to the combined genomic index containing both human and *P. falciparum* genomes using the splice-aware STAR aligner, allowing up to 8 mismatches for each paired-end read[52]. Reads were extracted from the output BAM file to separate parasite-mapped reads from human-mapped reads. Reads mapping to both genomes were counted for each sample and removed. BAM files were sorted, read groups replaced with a single new read group and all reads assigned to it, and indexed to run RNA-SeQC, a tool for computing quality control metrics for RNA-seq data[53]. HTSeq-count was used to count the reads mapped to exons with the parameter “-m union”[54]. Only uniquely mapping reads were counted.

Since our analysis of *P. falciparum* gene expression was reliant on a reference genome, families of highly polymorphic *var*, *stevor*, and *rifin* genes were removed from downstream analyses as these exhibit great sequence diversity between parasites and are likely to be incorrectly characterized[12]. Additional highly polymorphic regions within the *P. falciparum* genome which might also be incorrectly characterized were identified using schizont stage RNA-seq data from 9 clinical isolates (Duffy et al., manuscript submitted). In total, 139 genes were identified with highly polymorphic regions. A reference GTF file containing *P. falciparum* gene annotations was modified to remove these regions without removing the genes, and the resulting read count data generated using the modified GTF file was used for downstream analysis.

### Outlier identification

With the R package edgeR, raw read counts of each data set were normalized using a trimmed mean of M-values (TMM), which takes into account the library size and the RNA composition of the input data[55]. A multi-dimensional scaling (MDS) plot was used to identify the distances between samples that correspond to leading biological coefficient of variation. Up to the 6th dimension of MDS was plotted to fully observe the variation between samples, with two dimensions visualized at a time in scatter plot format. Three parasite samples were consistently found to be positioned away from other samples in each pair of dimensions, indicating outliers. This was further supported by low correlations observed between either of these outliers with other samples. These three samples were excluded from further parasite gene expression analysis: one sample (HL_478) had very low parasite reads making estimation of gene expression impossible and the other two samples (CH_285 and UM_589) were conspicuous outliers on MDS plots, possibly due to imperfect library preparation.

### Deconvolution analysis

To account for inter-individual variation in the proportions of different types of blood leukocyte, and for variation in the distribution of circulating parasites through the intraerythrocytic developmental cycle, deconvolution analysis was performed on RNA-seq data using CellCODE[15]. This uses a multi-step statistical framework to compute the relative differences in cell proportion represented as surrogate proportion variables (SPVs). It requires a reference data set that contains gene expression profiles for each cell type of interest. Five major immune cell populations were selected from Immune Response In Silico (IRIS)[56] to constitute the human reference data set: neutrophil, monocyte, CD4+ T-cell, CD8+ T-cell, and B-cell. Fragments Per Kilobase of transcript per Million mapped reads (FPKM) values were calculated from human RNA-seq data and log-transformed to simulate a microarray data set. For the parasite reference data set, RNA-seq data sets were obtained for four specific stages in the parasite asexual and sexual stage (0 hour, 24 hour, 48 hour, and gametocyte stage V)[16, 18], normalized by relative library sizes of samples (i.e. size factors) using edgeR. An identical normalization method was also applied for the input parasite RNA-seq data. A trial-and-error approach was taken to obtain the optimum SPV values for each cell-type. For human deconvolution, a cutoff value of 1.2 and a maximum number of marker genes of 50 appeared optimal. For parasite deconvolution, a cutoff value of 1.7 and a maximum number of marker genes of 50 appeared optimal.

Validation of CellCODE for *P. falciparum* developmental stage deconvolution (for which its use has not previously been reported) was performed by comparison with previously reported “stage-specific” marker genes[57] and by assessing performance in synthetic data sets constructed by mixing together in varying proportions randomly selected reads from RNA-seq reference datasets[16, 18] of the different parasite developmental stages.

### Differential gene expression and linear regression analysis

Prior to carrying out any downstream analyses genes with very low TMM-normalized read counts (< 5 counts-per-million (cpm) in < 3 samples and undetected in the remainder) were excluded. The generalized linear model tool in edgeR was employed to perform differential gene expression analysis (DGEA) between disease groups with adjustment for leukocyte and parasite SPVs, and in subsequent analyses additional adjustment for log parasite density and log PfHRP2.

Linear regression analysis was performed in edgeR to identify genes significantly associated with clinical variables of interest. Input gene expression values included adjustment for SPVs. The variables considered were: log PfHRP2, log parasite density, lactate concentration, platelet counts, and hemoglobin concentration. Additional analysis for lactate, platelets and hemoglobin were conducted including adjustment for log parasite density and log PfHRP2.

In both DGEA and linear regression analyses, false Discovery Rate (FDR) was computed for each individual analysis using the Benjamini-Hochberg procedure[58]. Genes with FDR below 0.05 were considered to be differentially expressed.

### Gene ontology and KEGG pathway enrichment analysis

Gene ontology (GO) terms for genes were obtained from Bioconductor package “org.Hs.eg.db” for human and “org.Pf.plasmo.db” for parasite. Input gene lists were significantly differentially expressed genes or genes that were significantly associated with laboratory variables. Fisher’s exact test was used to identify significantly over-represented GO terms from these gene lists. The background sets for each species consisted of all expressed genes detected in the data set with the exclusion of those with very low expression as described above. Enrichment analysis for biological process terms was carried out using the “goana()” function in edgeR. The least redundant GO terms with greatest significance in each analysis were identified for reporting using the tool REVIGO[59].

Ingenuity Pathway Analysis (Qiagen) was used for prediction of upstream regulators of groups of differentially expressed genes, and to identify functional networks.

### Construction of a coexpression network

The weighted gene coexpression network analysis (WGCNA) tool was used to construct a gene coexpression network[38]. The input data for WGCNA was read counts for each gene feature normalized using TMM method and then adjusted for SPVs using the command “removeBatchEffect()” from the R package edgeR. Both human and parasite expression data were analyzed together as a single set of genes for each subject. In order to comprehensively study the relationships between genes, two sets of networks were created: one with all samples from SM and UM groups, and the other with two separate sub-networks, generated from samples from SM and UM groups respectively. Network creation was conducted following the WGCNA tool guidelines:

1. Hierarchical clustering was performed at a sample level to detect outliers based on the WGCNA tool threshold, which were removed from the subsequent network generation (HL_171 and UM_492).
2. An appropriate soft-thresholding power (b) was chosen by applying the scale-free topology criterion. This was such that the power value enables the resulting gene network to satisfy the scale-free topology of approximately (*R*^2^ > 0.80).
3. Adjacency, which represents the connection strength of two genes in a network, was calculated. Coexpression similarity was calculated by taking the absolute value of the correlation coefficient, multiplying by 0.5 and adding 0.5 to create a signed network, where the presence of strongly negatively correlated gene pairs is downsized.
4. The adjacency matrix was transformed into a topological overlap matrix (TOM) in order to minimize the effects of spurious associations and noise in the network.
5. Hierarchical clustering on TOM dissimilarity was done to create hierarchical clustering tree of genes.
6. The dynamic tree cut method was used to group the genes that are highly correlated with one another into gene modules where minimum module size and the tree height at which genes below the height is grouped together were specified.
7. Module eigengene values for each module were calculated, which represents the overall gene expression profile of a module. Correlation analysis between modules was performed using eigengene values to identify modules with high similarity, which were then merged together.

The resulting network consisted of genes (represented as nodes in the network) and correlations between genes (represented as edges in the network), and highly correlated genes grouped together into modules. To characterize the gene network, several analysis steps were carried out. The most connected genes in each module were identified as the hub genes. Based on the module eigengene value, the connections between modules were determined. Pearson correlation analysis between module eigengene values and clinical variables was performed to identify gene clusters that are highly associated with clinical traits. Gene set enrichment analysis was performed on each module to identify significantly enriched GO terms. This data was summarised using OmicCircos[60].

From two separate sub-networks generated from SM and UM groups respectively, the preservation of gene connections across SM and UM groups was determined by assessing an overlap of genes for each module pair (from SM and UM sub-networks respectively), the significance of overlap was measured using the hypergeometric test. For each significantly preserved module pair, the hub genes and the significantly enriched GO terms were compared.

The gene network was exported to Cytoscape (<u>http://www.cytoscape.org/</u>) for visualisation. Only gene pairs with adjacency value of 0.03 or higher were exported to remove genes with low connections from the network visualization. SM and UM sub-networks were exported separately and subsequently combined into a single network using the Cytoscape embedded tool “Merge". By doing so, duplicate genes representing overlap between SM and UM sub-networks were removed, and connections between genes remained intact such that genes that can only be found on SM sub-network and also connected to the genes that can be found on both networks were not connected to the genes that can be found on UM sub-network and also connected to the same overlapping genes.

### Logistic regression for association of module eigengenes with severity

Logistic regression was performed using the glm package in R to identify module eigengene values with univariate association with severity. All modules with significant univariate associations (P<0.01) in addition to log PfHRP2 concentration were used in backward selection to identify the best multivariate model in which all terms were significant.

## Data availability

Sequence data that support the findings of this study will be deposited in ArrayExpress with the accession codes made available at the time of publication. Source data for Supplementary Table 1 are provided with the paper as Supplementary Dataset 1.

## Acknowledgements

This work was funded by the UK Medical Research Council (MRC) and the UK Department for International Development (DFID) under the MRC/DFID Concordat agreement and is also part of the EDCTP2 programme supported by the European Union (MR/L006529/1), MRC core funding of the MRC Gambia Unit (MRCG), and Wellcome Trust (098051). We are grateful to the study subjects, staff at MRCG, Jammeh Foundation for Peace Hospital, and Brikama Health Centre; Konrad Paszkiewicz and staff at Exeter Sequencing Service at the University of Exeter (supported by Medical Research Council Clinical Infrastructure award (MR/M008924/1), Wellcome Trust Institutional Strategic Support Fund (WT097835MF), Wellcome Trust Multi User Equipment Award (WT101650MA) and BBSRC LOLA award (BB/K003240/1)).

## Author contributions

AJC, ML and DJC conceived the study; HJL, TDO and AG performed formal analysis; MW and AJC performed investigations on clinical samples; MW, DJC, DN and LBS provided samples; DJC, TDO and LBS provided methodology; AJC and HJL wrote the original draft; all authors contributed to review and editing of the manuscript; LJC, DJC, ML, TDO and AJC provided supervision; AJC obtained funding.

## Materials & Correspondence

Correspondence should be addressed to Dr Aubrey Cunnington, Clinical Senior Lecturer, Section of Paediatrics, Department of Medicine, Imperial College London (St Mary’s Campus) 231, Medical School Building, Norfolk Place, London W2 1PG.

## Supplementary Material

**Supplementary Table 1.** Characteristics of study subjects (n=46)

|  | CM (n=5) | CH (n=12) | HL (n=8) | UM (n=21) | *P* (*F* value) |
| --- | --- | --- | --- | --- | --- |
| **Age (years)** | 4.3 (4.2-4.8) | 4.9 (3.6-5.7) | 5.0 (3.8-8.3) | 6.0 (4.0-9.0) | 0.24<br>(1.47) |
| **Male (%)** | 3 (60%) | 5 (42%) | 7 (88%) | 13 (62%) | 0.24 |
| **Parasitemia (%)** | 8.3 (5.3-9.0)<sup>4</sup> | 12.6 (9.4-19.0) | 9.6 (1.8-12.2) | 5.1 (3.8-7.0) | 0.02<br>(3.67) |
| **Parasites (x10<sup>5</sup>/uL)** | 2.3 (1.7-3.1)<sup>3</sup> | 3.5 (2.7-8.4)<sup>11</sup> | 2.8 (0.7-5.0) | 2.3 (1.6-3.2) | 0.02<br>(3.55) |
| **Clones** | 2 (1.5-2.5)<sup>4</sup> | 2 (1-2)<sup>9</sup> | 1 (1-2)<sup>5</sup> | 2 (1-2)<sup>15</sup> | 0.68<br>(0.51) |
| ***Pf*HRP2 (ng/mL)** | 202 (93-528)<sup>4</sup> | 763 (374-1750) | 470 (164-2214) | 163 (128-227) | 0.001<br>(6.20) |
| **Duration of illness (days)** | 2.0 (1.7-3.0) | 2.0 (2.0-2.5) | 2.0 (2.0-3.5) | 2.7 (2.0-3.0) | 0.57<br>(0.68) |
| **Hb (g/dL)** | 9.7 (7.4-10.4) | 9.3 (7.8-11.5)<sup>11</sup> | 9.1 (7.4-11.0) | 10.8 (9.9-12.1) | 0.10<br>(2.24) |
| **WBC (x10<sup>9</sup>/L)** | 9.8 (8.2-12.9)<sup>4</sup> | 8.8 (6.4-9.4)<sup>11</sup> | 15.3 (7.9-16.8)<sup>7</sup> | 9.5 (7.7-11.8) | 0.30<br>(1.27) |
| **Platelets (x10<sup>9</sup>/L)** | 41 (40-82)<sup>4</sup> | 36 (23-65)<sup>11</sup> | 59 (33-132) | 122 (96-132) | 0.04<br>(3.15) |
| **Lymphocyte (%)** | 29.8<br>(20.6-37.3)<sup>4</sup> | 37.8<br>(29.9-49.9)<sup>11</sup> | 22.3<br>(14.7-37.3) | 23.9<br>(16.0-33.5)<sup>20</sup> | 0.05<br>(2.78) |
| **Neutrophil (%)** | 55.1<br>(49.0-69.6)<sup>4</sup> | 48.3<br>(39.6-56.2)<sup>10</sup> | 61.5<br>(55.6-74.9)<sup>7</sup> | 68.0<br>(59.9-79.6)<sup>20</sup> | 0.01<br>(4.25) |
| **Monocyte (%)** | 7.1<br>(6.0-7.7)<sup>4</sup> | 7.8<br>(6.8-8.6)<sup>10</sup> | 6.6<br>(5.1-7.8)<sup>7</sup> | 6.7<br>(4.8-7.3)<sup>20</sup> | 0.38<br>(1.05) |

CM, cerebral malaria; CH, cerebral malaria plus hyperlactatemia; HL, hyperlactatemia (CM, CH, and HL are all subgroups of severe malaria, SM); UM, uncomplicated malaria. Data are median (IQR), superscripts indicate the number of subjects with data for each variable if less than the total; *P* (*F*) for ANOVA comparing all groups (degrees of freedom =3) except for sex where *P* is for Fisher's exact test.

**Supplementary Table 2. Human genes differentially expressed between severe malaria phenotypes and uncomplicated malaria in unadjusted and parasite load-adjusted analyses.**

**Supplementary Table 3. *P. falciparum* genes differentially expressed between severe malaria phenotypes and uncomplicated malaria.**

**Supplementary Table 4. Human genes significantly correlated with parasite load measurements and laboratory parameters.**

**Supplementary Table 5. *P.falciparum* genes significantly correlated with parasite load and laboratory parameters.**

**Supplementary Table 6. Gene ontology terms associated with human differentially expressed or significantly correlated genes in unadjusted and parasite load-adjusted analyses.**

**Supplementary Table 7. Predicted upstream regulators associated with human differentially expressed or significantly correlated genes in unadjusted and parasite load-adjusted analyses.**

**Supplementary Table 8. Gene ontology terms associated with parasite differentially expressed or significantly correlated genes in unadjusted and parasite load-adjusted analyses.**

**Supplementary Table 9. Summary of modules obtained from combined whole genome correlation network.**

**Supplementary Table 10.** Univariate and multivariate associations of module eigengene values and parasite load with severity.

|  | Univariate<br>log odds | Univariate<br>P-value | Multivariate<br>log odds | Multivariate<br>P-value |
| --- | --- | --- | --- | --- |
| **MMP8** | 12.4 | 0.00130 | **66.1** | **0.0152** |
| **LYSMD3** | 9.49 | 0.00353 | **28.6** | **0.0178** |
| PF3D7_1129400 | 10.0 | 0.00388 |  |  |
| RPL24 | 7.77 | 0.00463 |  |  |
| **OAS3** | -6.73 | 0.00943 | **47.7** | **0.0261** |
| HSPH1 | 9.74 | 0.0117 |  |  |
| TNRC6B | -6.44 | 0.0136 |  |  |
| KIF11 | 6.15 | 0.0179 |  |  |
| PF3D7_1415300 | -8.77 | 0.0184 |  |  |
| PF3D7_1252500 | -5.50 | 0.0216 |  |  |
| PF3D7_1105600 | -5.91 | 0.0218 |  |  |
| LMAN1 | 5.63 | 0.0255 |  |  |
| PF3D7_0721600 | 5.33 | 0.0283 |  |  |
| PF3D7_0520100 | 4.99 | 0.0307 |  |  |
| TPM3 | 4.44 | 0.0560 |  |  |
| ANKRD49 | 3.43 | 0.133 |  |  |
| MT.ND4 | 2.48 | 0.251 |  |  |
| PF3D7_0405400 | 2.18 | 0.289 |  |  |
| MUC19 | -2.18 | 0.315 |  |  |
| PF3D7_0812900 | -1.85 | 0.367 |  |  |
| SH3BGRL2 | -1.41 | 0.491 |  |  |
| PSMC3 | 1.17 | 0.568 |  |  |
| PF3D7_1024800 | -1.08 | 0.596 |  |  |
| PF3D7_1138700 | 0.938 | 0.643 |  |  |
| CPSF6 | 0.647402239 | 0.747 |  |  |
| MPP1 | 0.02183521 | 0.991 |  |  |
| Log parasite density | 1.17 | 0.0339 |  |  |
| Log PfHRP2 | 1.4031 | 0.00440 |  |  |

Log odds are per unit change in the variable, calculated using logistic regression. The multivariate model was derived by backwards selection from all variables with univariate $P < 0.01$ .

**Supplementary Table 11. Summary and overlap of whole genome correlation sub networks for severe and uncomplicated malaria**

**Supplementary Fig 1.**
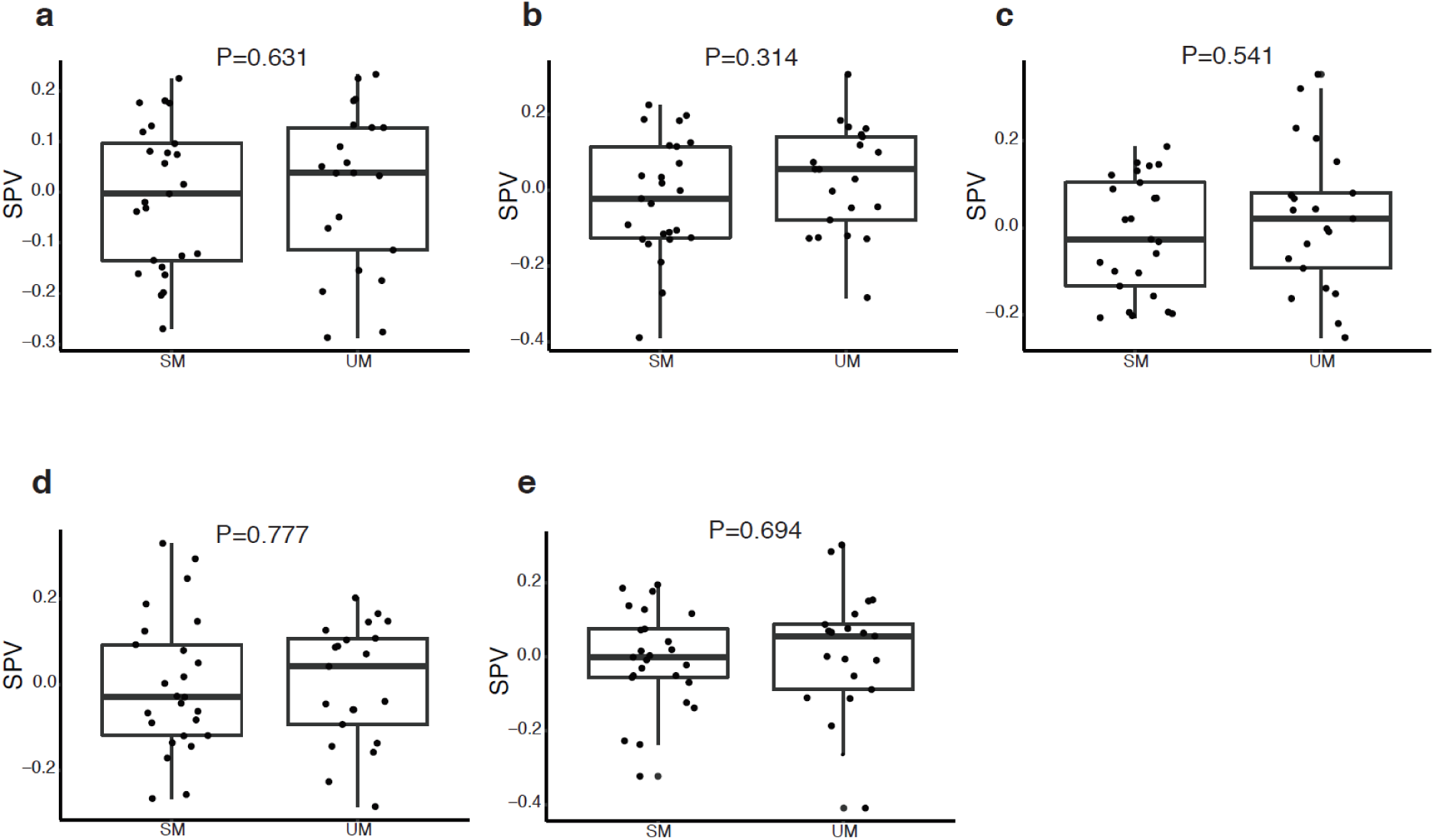
Estimates of the relative proportions of leukocyte subpopulations in subjects with severe and uncomplicated malaria. **(a-e)** surrogate proportion variables compared by severity category for neutrophils **(a)**, monocytes **(b)**, CD4+ T – lymphocytes **(c)**, CD8+ T – lymphocytes **(d)**, and B-lymphocytes **(e)** using the Mann – Whitney test (UM, n=21; SM, n=25; boldline, box and whiskers indicate median, interquartile range respectively)

**Supplementary Fig 2.**
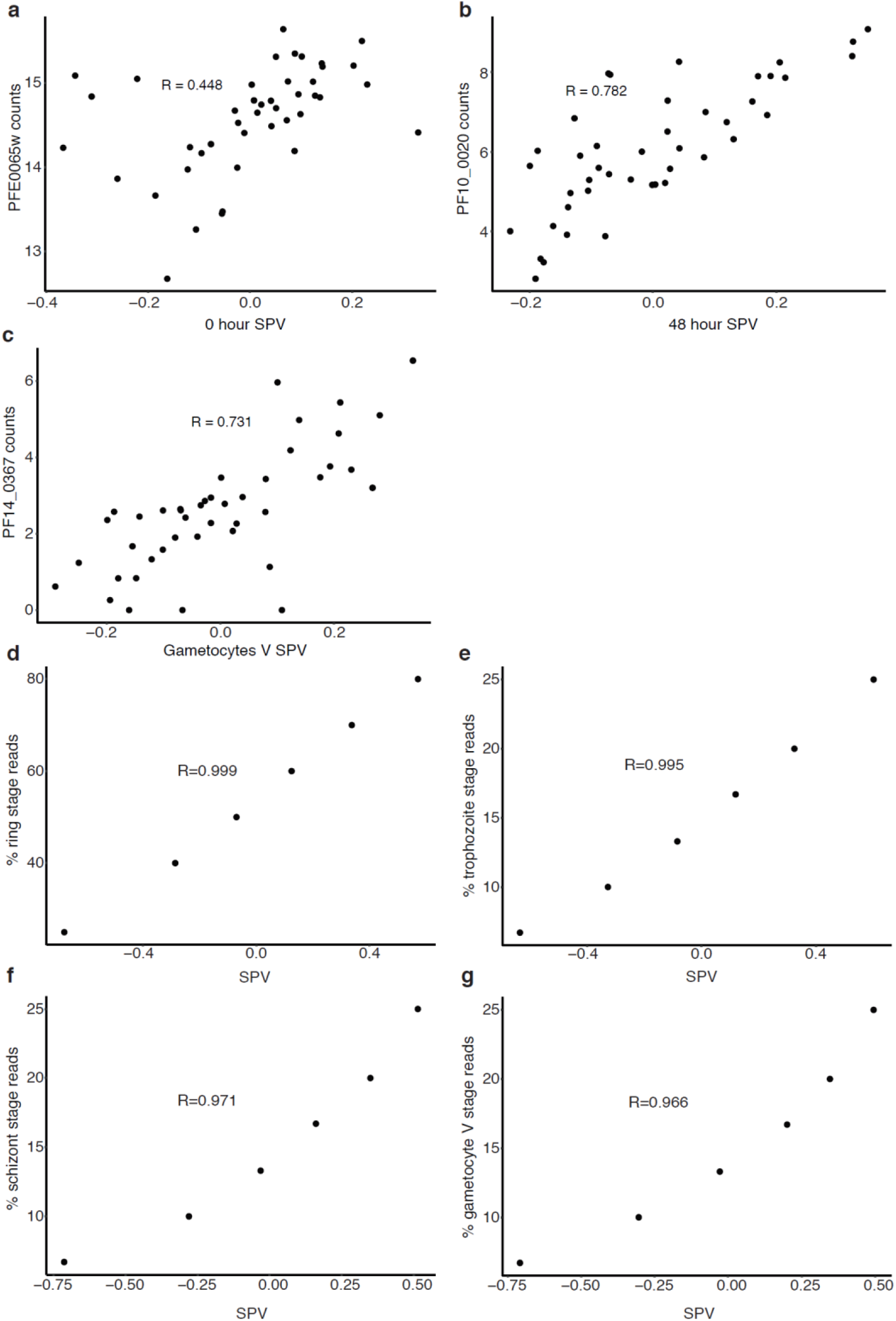
Validation of the gene – signature approach to estimate parasite developmental satage proportions (a-c) Correlation of surrogate proportion variables (SPV) with read counts for putataive “stage – specific” marker genes (see Methods): **(a)** 0hr SPV vs early asexual stage marker gene PFE0065w; **(b)** 48hr SPV vs late asexual stage marker gene PF10_0020; **(c)** Gametocyte V SPV vs developing gametocyte marker gene PF14_0367 (R for pearson correlation, n=43). (d-g) Correlation of SPVs with actual proportion of reads derived from each parasite developmenatal stage in synthetic mixtures of varying proportions of stage – specific RNA – seq reads from early ring – stage (0 hour, **d**), trophozoite (24 hour, **e**), late schizont (48 hour, **f**) and mature gametocyte (stage v, **g**) R for pearson correlation, n=43.

**Supplementary Fig 3.**
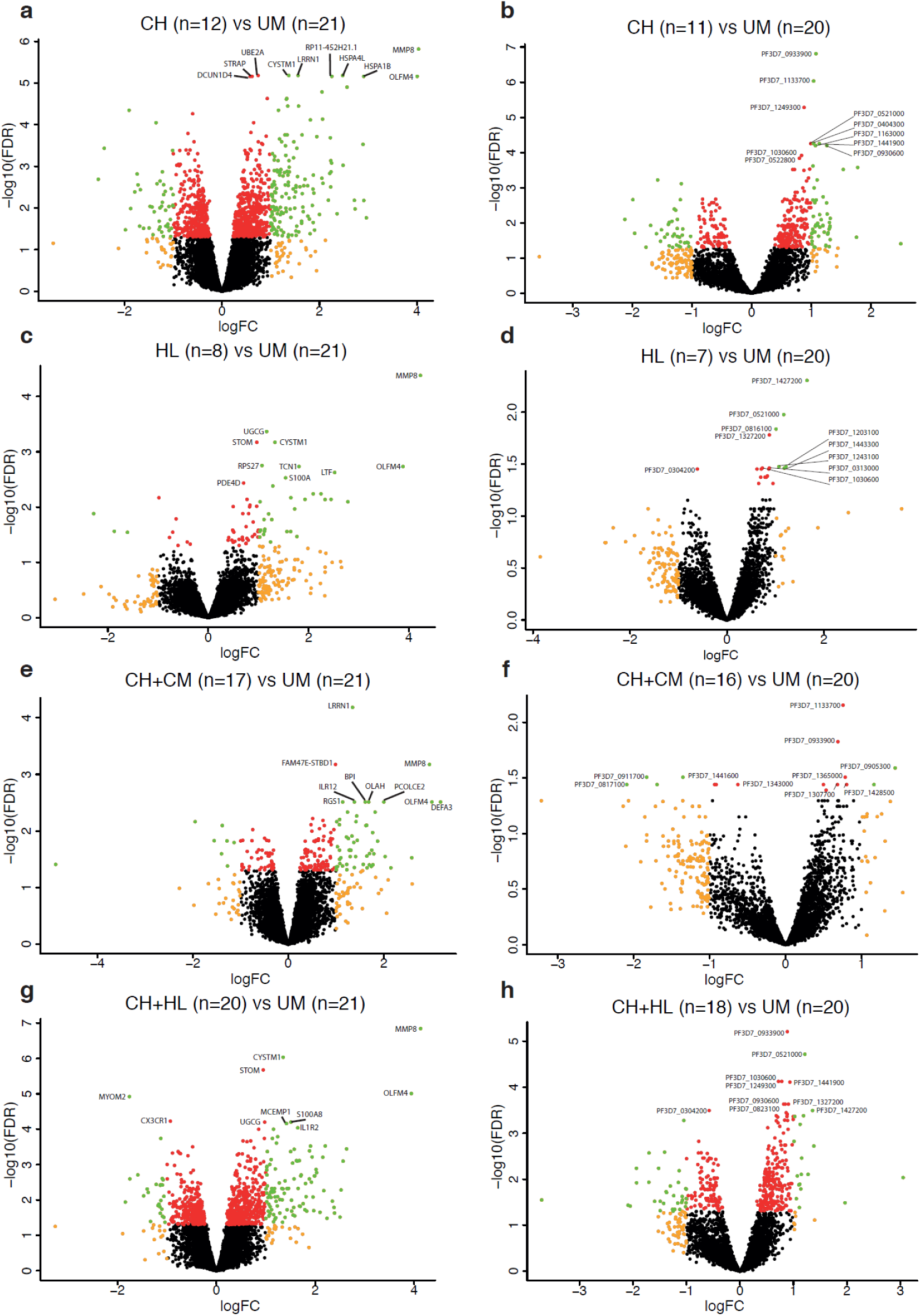
Differential gene expression between severe malaria phenotypes and uncomplicated malaria. Volcano plots showing extent and significance of up- or down- regulation of human (left hand column) or *P. falciparum* (right hand column) gene expression in comparisons between specific phenotypes of SM vs UM (red and green, P <0.05 after Benjamini-Hochberg adjustment for false discovery rate (FDR); orange and green, absolute log_2_-fold change (FC) in expression > 1; the 10 most significant genes are annotated).

**Supplementary Fig 4.**
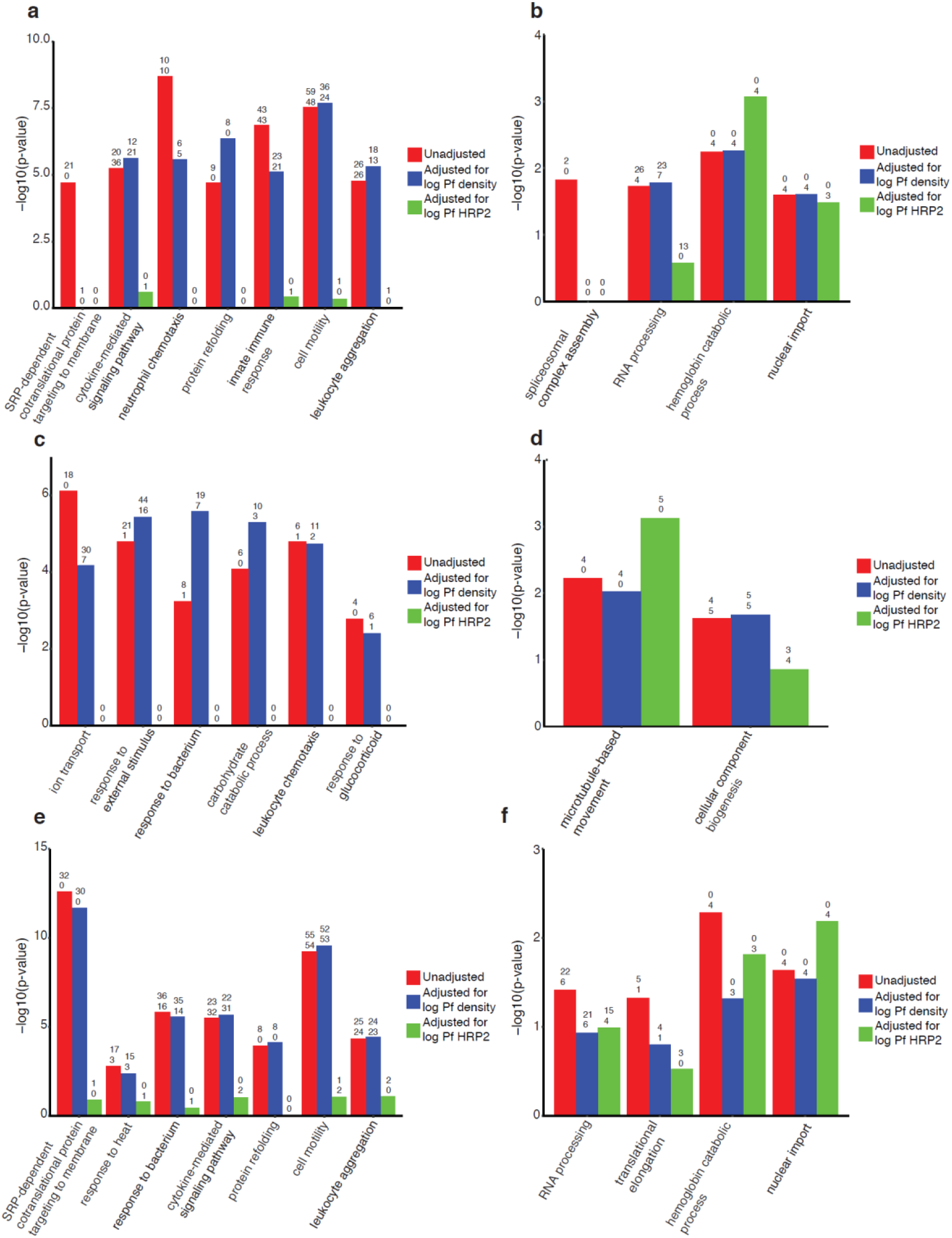
Gene ontology terms associated with genes differentially expressed between severe malaria phenotypes and uncomplicated malaria, unadjusted or adjusted for parasite load. Most significantly enriched, non-redundant, gene ontology terms for genes significantly differentially expressed (DEGs) between SM phenotypes and UM (numbers above each bar indicate the number of up- regulated and down-regulated genes within each category). Comparisons are only shown if they include multiple significantly enriched GO terms. (**a**) CH (n=11) vs UM (n=21) human DEGS, (**b**) CH (n=11) vs UM (n=20) *P. falciparum* DEGs, (**c**) HL (n=8) vs UM (n=21) human DEGS,(**d**) CH+CM (n=14) vs UM (n=20) *P. falciparum* DEGs, (**e**) CH+HL (n=19) vs UM (n=21) human DEGS, (**f**) CH+HL (n=18) vs UM (n=20) *P. falciparum* DEGs.

**Supplementary Fig 5.**
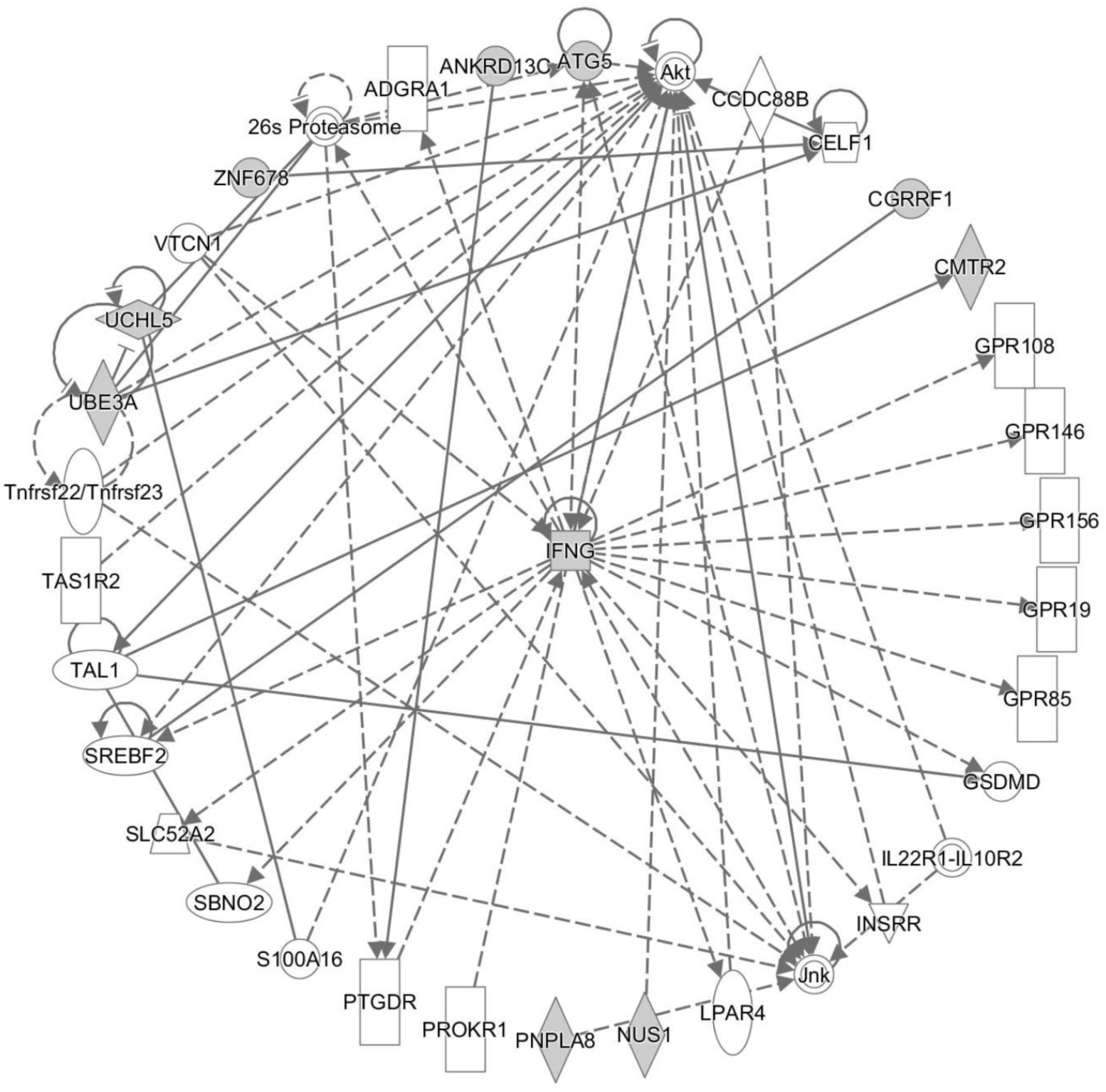
Top functional network for the small LYSMD3 module. Functional networks were identified in Ingenuity Pathway Analysis software and the top scoring network is portrayed in radial layout which places the most interconnected gene at the centre. Genes within the module are shaded.

**Supplementary Dataset 1. Subject-level clinical and laboratory data**

